# Scalable genomics: from raw data to aligned reads on Apache YARN

**DOI:** 10.1101/071092

**Authors:** Francesco Versaci, Luca Pireddu, Gianluigi Zanetti

## Abstract

The adoption of Big Data technologies can potentially boost the scalability of data-driven biology and health workflows by orders of magnitude. Consider, for instance, that technologies in the Hadoop ecosystem have been successfully used in data-driven industry to scale their processes to levels much larger than any biological- or health-driven work attempted thus far. In this work we demonstrate the scalability of a sequence alignment pipeline based on technologies from the Hadoop ecosystem – namely, Apache Flink and Hadoop MapReduce, both running on the distributed Apache YARN platform. Unlike previous work, our pipeline starts processing directly from the raw BCL data produced by Illumina sequencers. A Flink-based distributed algorithm reconstructs reads from the Illumina BCL data, and then demultiplexes them – analogously to the bcl2fastq2 program provided by Illumina. Subsequently, the BWA-MEM-based distributed aligner from the Seal project is used to perform read mapping on the YARN platform. While the standard programs by Illumina and BWA-MEM are limited to shared-memory parallelism (multi-threading), our solution is completely distributed and can scale across a large number of computing nodes. Results show excellent pipeline scalability, linear in the number of nodes. In addition, this approach automatically benefits from the robustness to hardware failure and transient cluster problems provided by the YARN platform, as well as the scalability of the Hadoop Distributed File System. Moreover, this YARN-based approach complements the up-and-coming version 4 of the GATK toolkit, which is based on Spark and therefore can run on YARN. Together, they can be used to form a scalable complete YARN-based variant calling pipeline for Illumina data, which will be further improved with the arrival of distributed in-memory filesystem technology such as Apache Arrow, thus removing the need to write intermediate data to disk.

**Original article:** This paper was presented at the IEEE International Conference on Big Data, 2016 and is available at https://doi.org/10.1109/BigData.2016.7840727

## I. Introduction

The data-intensive revolution in the life sciences [1], [2] has next-generation sequencing (NGS) machines among its most prominent officers. As it becomes more economically accessible, DNA sequencing has opened up to a myriad of new applications that were previously technologically or economically unfeasible [3]. High-throughput sequencing can now be used for research into understanding human genetic diseases [4], in oncology [5], to study human phylogeny [6], and is even reaching the level of personalized diagnostic applications [7].

One of the main challenges brought forth by this phenomenon is to develop scalable computing tools that can keep up with such a massive data generation throughput. The raw data produced by NGS needs to go through various intense processing steps to extract biologically relevant information. To date, it appears that most sequencing centers have opted to implement processing systems based on conventional software running on High-Performance Computing (HPC) infrastructure [8] – a set of computing nodes accessed through a batch queuing system and equipped with a parallel shared storage system. While with enough effort and equipment this solution can certainly be made to work, it presents some issues that need to be addressed. Two important ones are that developers need to implement a general way to divide the work of a single job among all computing nodes and, since the probability of node failures increases with the number of nodes, they also need to make the system robust to transient or permanent hardware failures, recovering automatically and bringing the job to successful completion. Nevertheless, even with these measures, the architecture of the HPC cluster limits the maximum throughput of the system because it is, usually, centered around a single shared storage volume, which tends to become the bottleneck as the number of computing nodes increases – and this is especially true for some phases of sequence processing which can perform a lot of I/O with respect to processing activity.

This paper presents our novel approach to processing sequencing data, adopting a strategy completely different from the status quo by processing raw sequencing data using Hadoop MapReduce and Apache Flink (see Sections III-A and III-B). To the best of the authors’ knowledge, this is the first solution that can process the sequencer’s raw data directly on a distributed platform. In brief, in this work we present:

- the first complete and scalable Hadoop-based pipeline to align DNA sequences starting from raw data;
- an efficient Flink-based tool to convert from BCL to FASTQ formats;
- the Read Aligner API (RAPI), which encapsulates aligner functionality and provides C and Python bindings;
- improvements to the efficiency of the aligner in the Seal suite.

The rest of the paper is organized as follows. Section II describes the Next Generation Sequencing process, the computation that is required to make sense of the data and the state-of-the-art in modern sequencing centers. Section III provides background regarding Hadoop and Flink, motivating the decision to adopt the Hadoop/YARN framework. Sections IV and V present the tools that have been developed as part of this work. Section VI contains the performance evaluation and a comparison to the state-of-the-art. Finally, Section VIII discusses related work and Section IX concludes the manuscript.

## II. The NGS process

Deoxyribonucleic acid (DNA) is a polymer composed of simpler units known as nucleotides, or *bases*. These come in four kinds: Adenine, Cytosine, Guanine and Thymine – respectively denoted by their initial letters A, C, G, and T. The DNA data produced by the NGS process are not directly interpretable from a biological point of view. In fact, the various high-throughput NGS technologies [9] all use a “shotgun” approach, where the genome to be sequenced is broken up into fragments of approximately the same size and the individual fragments are sequenced in parallel. The characteristics of the raw data produced by the sequencer changes depending on the specific technology being used. The sequencers by Illumina Inc. (http://www.illumina.com) – which are the target of this work – operate by successively attaching a fluorescent identifying molecule to each base of the DNA fragments being sequenced [10].

Thus, at each cycle, the machine acquires a single base from all the reads by snapping a picture of the flowcell where a chemiluminescent reaction is taking place. For each picture, and thus cycle, the software on the sequencer’s controlling workstation performs base calling from the image data – mapping each chemiluminescent dot to a specific base (A, C, G, T) based on its color. The process produces base call files (BCL), which contain the bases that were acquired from all the fragments – also known as *reads* – but only in that specific sequencing cycle. Therefore, the individual DNA fragments are actually split over *C* files, where *C* is the number of cycles in the sequencing run and the length of the reads.

Next-generation sequencing activity quickly results in a lot of data. Consider that a modern high-throughput sequencer like the Illumina HiSeq 4000 can produce 1500 Gigabases, which equate to about 4 Terabytes of uncompressed sequencing data, in just 3.5 days [11]. That much data is sufficient to reconstruct up to 12 human-sized genomes, where a single sample of this type equates to a bit over 300 GB of uncompressed data. Moreover, in the context of a study or a sequencing center, this hefty per-sample data size is typically multiplied by a significant number of samples. Consider that for sequencing-based studies that require high analysis sensitivity or to get population-wide statistics, thousands of samples need to be sequenced and processed; e.g., a population-wide study by Orrù et al. [4] required the sequencing of 2870 whole genomes resulting in almost one petabyte of sequencing data to be analyzed and stored.

**TABLE I:** Maximum sequencing capacities for a number of Illumina high-throughput sequencers [11]–[13]. The size of the output in GB is intended for uncompressed reads with base qualities and id strings, considering a total of 2.5 bytes/base.

| Sequencer | Read length | Time (days) | Gigabases | Gigabytes (GB) | Rate (GB/day) |
| --- | --- | --- | --- | --- | --- |
| HiSeq 2000 | 2 x 100 | 8 | 400 | 1000 | 125 |
| HiSeq 2500 | 2 x 125 | 6 | 1000 | 2500 | 417 |
| HiSeq 4000 | 2 x 150 | 3.5 | 1500 | 3750 | 1071 |

### A. BCL conversion and sorting reads

The first step in making sense of the raw data is to reconstruct the original DNA fragments from the “slices” produced by the sequencer. The operation required to reconstruct the reads is logically equivalent to a matrix transposition, where a read is obtained by concatenating the elements located at the same positions across several files, as illustrated in Fig. 1.

**Fig. 1:**
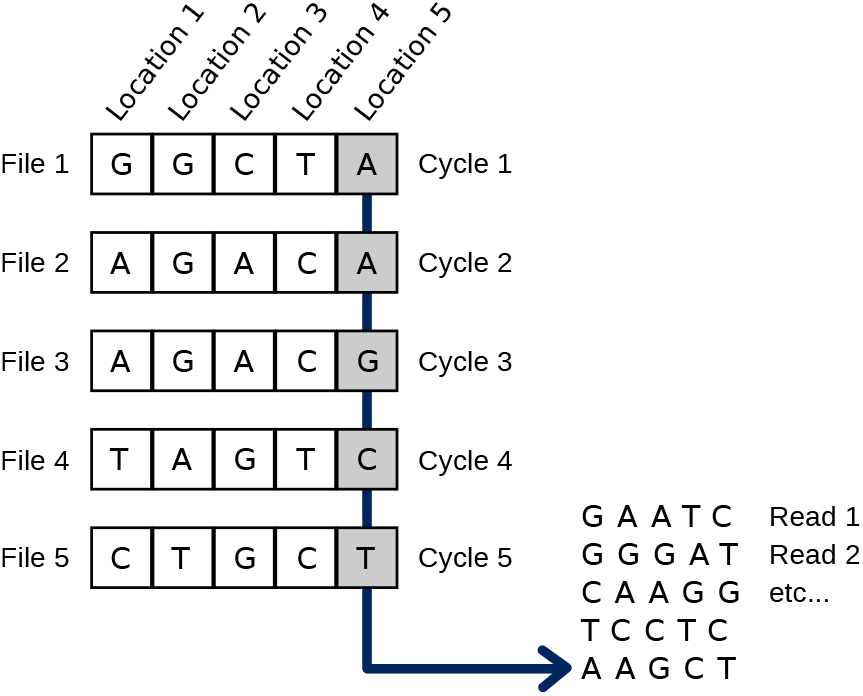
Recomposing nucleotide sequences from the raw data in Illumina BCL files. The operation requires reading data simultaneously from a number of files equal to the number of cycles in the sequencing run – each file contains one base from each read. Once all the bases of a read are acquired, the read can be composed and emitted and processing can advance to the next one.

Along with the read reconstruction, one must typically also perform other operations which help the subsequent analyses: filtering out some of the reads (based on filter files which are also part of the run output) and tagging each read with some metadata (e.g., location of read within the flowcell).

Moreover, for reasons of efficiency and flexibility, the DNA fragments from individual samples are often tagged with a short identifying nucleotide sequence and then mixed and sequenced with other samples in a single batch. This procedure is known as *multiplexed sequencing*. In this way, the sequencing capacity of the run is divided among more biological samples than would be possible if it was necessary to keep them physically separate. Being that the genetic material of multiple samples is in the same biological solution, the sequencer will acquire their fragments of DNA indiscriminately and they will be output in the same dataset. Thus, after BCL conversion, the NGS data will typically undergo a *demultiplexing* stage, where fragments from various samples are sorted into separate datasets using their identifying tag.

### B. Read Alignment

The product of the high-throughput shotgun sequencing and processing procedure described thus far is a set of billions unordered short DNA fragments. These need to be analyzed computationally to understand the structure of the original whole DNA sequence from which they came. The specific analyses to be performed will vary depending on the application. The most common sequencing scenario – and the one that concerns us in this work – is the one where there is a reference genome available for the sample being sequenced and it is used as a guide to reconstruct the sample’s genome. This type of sequencing experiment is known as *resequencing.* When resequencing, after the reconstruction of the sequenced fragments, the fragments are *aligned* or *mapped* to the reference genome – i.e., an approximate string matching algorithm is applied to find the position in the reference sequence from which the reads were most likely acquired. The mapping algorithms are designed to be robust to sequencing errors and to the small genetic variations that characterize the sample. Moreover, when sequenced the genome is typically oversampled many times; for instance, for whole-genome sequencing 30 times oversampling – also known as *30X coverage* is common. The oversampling results in reads that overlap, which will be essential in later phases of analysis to detect heterozygosity and to distinguish genomic variants specific to the sample from mere sequencing errors.

### C. Standard practice

In a typical scenario, after the sequencer finishes its work a multi-step processing pipeline will be executed, starting with the BCL conversion and read alignment steps described in the previous subsections. In the state-of-the-art, sequencing operations are equipped with a conventional HPC cluster with a shared parallel storage system. The nodes are typically accessed via a batch queuing system, and often the computing resources are dedicated to the needs of the sequencing and bioinformatics work [8].

Within this context, the standard solution is to perform read reconstruction and demultiplexing using Illumina’s own proprietary, open-source tool: bcl2fastq2. It is written in C++ and powered by the Boost library [14]. This tool implements shared-memory parallelism – i.e., it only exploits parallelism within a single computing node. To the best of the authors’ knowledge, there are no alternatives for this tool in a conventional computing setting.

On the other hand, there is variety of alignment programs available and in widespread use [15], [16]. Among these, BWA-MEM [17] is quite popular and has been found to produce some of the best alignments [18], [19]. Like the bcl2fastq2 – and the other conventional read alignment programs – BWA-MEM also implements shared-memory parallelism.

Though shared-memory parallelism is certainly beneficial to accelerating analysis, it is insufficient. Because of the huge amount of data produced by current sequencers the processing of a run on a single node easily takes several hours of computation, even on state-of-the-art machines, with tens or even hundreds of cores. One way to overcome this problem is to distribute the data among different nodes and then running distinct instances of the programs on each node, each one working on its subset of the data. However, this ad hoc solution is not trivial to implement properly, as one would inevitably end up trying to reimplement one of the already existing distributed computing frameworks.

## III. Background

### A. Hadoop framework

The Hadoop framework, which has its origins in data companies such as Yahoo! and Google, has been designed to scale computing throughput up to very high levels while containing costs. In particular it provides:

- Robustness to hardware failures, by automatically restarting tasks that do not complete, even because of a broken node, without any user intervention;
- A distributed file system (HDFS), which spreads data and allows them to be processed directly on the node where they are stored, drastically reducing the network load.

Another point worth mentioning is that by adopting Hadoop, one also acquires access to Hadoop-based Platform-as-a-Service (PaaS) offerings – for example, Amazon Elastic MapReduce (EMR) [20] Google Cloud Dataproc [21] – which entails having access to scalable cloud computing infrastructure without incurring any initial investment cost.

On the other hand Hadoop is not compatible with pre-existent software: to adopt this framework in a sequencing operation and reap the benefits it offers, one has to find or implement new Hadoop-compatible software to perform the processing steps required by the sequence processing workflow; this is the problem that the work presented here is addressing.

### B. Flink

Apache Flink [22] is a system for processing in parallel large streams of data; it is written in Java and can be run standalone or within an existent Hadoop/YARN installation [23], benefitting both from shared-and distributed-memory parallelism. It originates from the Stratosphere project [24], developed at TU Berlin, and is now a top-level project of the Apache Software Foundation, supported by a large community and being increasingly adopted both in research and industry.^1^

## IV. BCL processing

In this section we describe our BCL converter, which is shown to have the same performance as the shared-memory Illumina bcl2fastq2 tool on a single node, while also enabling easy and scalable distributed-memory parallelization. Our tool is written in Scala [25] within the Flink framework and can thus be easily integrated in any Hadoop/YARN system, as a simple module to be used in any Hadoop-based workflow.

The organization of the BCL files depends on the generating sequencer and in this paper we will adopt the format used by the HiSeq 3000/4000 machines. Files are divided into a tree structure, with a single file referring to data obtained by specific lane, tile and cycle combination: e.g., the file L003/C80.1/s_3_1213.bcl.gz corresponds to data read from tile *t* = 1213, in lane *l* = 3 during cycle *c* = 80. The uncompressed BCL files have constant size within a run and represent arrays of bytes *R*(*l,t,c*). Let’s now consider the matrix *M*(*l,t*), where the *c*-th row of *M*(*l*, *t*) is given by *R*(*l,t,c*). The first step of the BCL processing consists in computing the transposition of *M*(*l,t*), for all *l* and *t*. A typical run with 8 lanes, hundreds of tiles within each lane, hundreds of cycles per tile and BCL files of a few megabytes, produces a total output in the order of the terabyte, scattered among hundreds of thousands of files.

### A. Algorithmic details

From a high-level perspective our converter works as follows: the basic unit of job is the processing of a (lane, tile) combination and several of these jobs are grouped together into a Flink job, which is assigned a number of cores required to run. Flink jobs are scheduled by our program and sent to the Flink executor, which allocates them on the available resources. As an example, given a cluster of 8 nodes, with 16 cores/node and 2 threads/core a possible assignment of the resources is the following: 2 units of job per Flink job (to match the simultaneous multithreading), 1 core per Flink job, 128 Flink jobs running concurrently.

When processing a (lane, tile), paired-end sequences are handled at the same time, since they share the same *filter* and *locs* files, which contain respectively information to filter the sequence and to reconstruct the flowcell coordinates of each read. Furthermore, the reads will be used together during the alignment, so we have chosen to aggregate them already at this stage of the processing. The BCL files corresponding to different cycles are opened concurrently and the bases and quality scores are extracted and filtered (see IV-B for some implementation details). For each fragment, a header is added, containing various meta-data and the fragment’s multiplexing tag sequence, if present.

Once the fragments from the (lane, tile) have been reconstructed and tagged, they are sorted by their tag sequence. Since there can be read errors both in the data and in the tags, their repartition can be done in a fuzzy way: a parameter, which defaults to one, sets the numbers of allowed substitution errors when matching a tag (i.e., the Hamming distance between tag and match). Finally, a compressed file for each tag is written to disk, with files from different (lane, tile) combinations which match the same tag being written in the same directory (they belong to the same biological sample).

### B. Low-level optimizations

Our program is written in Scala and compiles into Java bytecode, so it has been challenging to match the performance of programs written in languages closer to the machine hardware. Nonetheless, our program matches, on the single computing node, the speed of bcl2fastq2 – i.e., Illumina’s proprietary tool. In order to achieve such high performance we used several techniques and optimizations, the most rewarding being presented below.

#### 1) Processing data in chunks

BCL files are arrays of bytes, in which each byte encodes a base (bits 0-1) and a quality score (bits 2-7). Therefore it would seem natural to process them in Flink as datastreams of bytes (i.e., using the `DataStream[Byte]` type). However, to exploit the full performance of the Flink engine we had to read and process data in bigger chunks (2048 bytes in our implementation), which better exploits cache locality and lowers the overhead of the streaming framework.

#### 2) Use of ByteBuffer

In order to decode the bases and quality scores encoded in the BCL files the bytes need to go through some bit masks and shifts. Since the operations to be performed are the same for every byte read, one can obtain a factor 8 speed-up by grouping bytes into 64-bit longs and executing the equivalent operations on longs. To implement this optimization in Scala we used the ByteBuffer class, which allows us to interpret byte arrays as longs.

#### 3) Use of look-up tables

We implemented the conversion of bases from numeric to ASCII notation (e.g., `0x0001020303020100` maps to “ACGTTGCA”) by “compressing” the input and using it as an index into a look-up table (e.g., `0x0001020303020100` is compressed to an index `0b0001101111100100=0x1BE4` into the precomputed look-up table).

## V. Aligning Reads

Read alignment in our read processing workflow was performed using Seal seqal – the Hadoop-based aligner in the Seal toolkit [26], [27] – which we improved as part of this work. Among the various Hadoop-based read aligners (see Section VIII) Seal was chosen because it integrates the BWA-MEM aligner, has good performance, and was written by some of the authors of this work.

The alignment step is implemented in seqal as a *map-only* job using Pydoop [28]. Rather than implementing a new aligner from scratch, seqal directly integrates the BWA-MEM aligner through the RAPI read aligner API (developed as part of this work; see Sec. V-A). To integrate BWA-MEM, its C source code was significantly modified to repackage it as a software library, so that its functions and data structures could be called by seqal. Indeed, currently seqal is, to the best of the authors knowledge, the only Hadoop-based read mapping program that directly integrates the aligner core into its code; other options choose to execute conventional read alignment programs in their unmodified form [29]–[33] as a separate child process. The technique adopted by seqal, while being more difficult to implement, provides improved flexibility and should also have a positive effect on efficiency.

In addition to transforming BWA-MEM into a software library, the original code was also modified to use memory-mapped I/O to read the reference sequence and its index, instead of simply writing the index components into `malloc’`d memory blocks. The reference index and sequence, to which the reads are mapped, is relatively large when working with higher order organisms; for instance, the human genome (3 Gigabases long) results in an indexed reference for BWA-MEM that is just over 5 GB in size. Memory mapping this data provides two significant advantages over BWA-MEM’s original technique. The first is that by using `mmap` all instances of the software using the same reference on the same computer will share the same data in memory, thereby significantly reducing the memory required for an alignment job. This feature is especially important to seqal’s use case since, for best efficiency, it should run many concurrent map tasks per node; empirically we have found that one task per hyperthreaded core achieves very good CPU utilization. The second advantage is that after its first use the reference is automatically cached in memory for a period of time, so that subsequent invocations of the aligner with the same reference will save the time required to load it from storage (a significant amount of time for larger references such as for *Homo sapiens*). Since the BWA-MEM project is open source, these improvements were sent to the author of the original program to be considered for integration into the main line of development.

### A. The RAPI Read Aligner API

In work related to NGS data, read aligners are currently one of the most fundamental pieces of analysis software, so read alignment is an operation that has been intensely studied [15], [34], resulting in a number of effective algorithms and implementations, many of which are continuously evolving.

These tools are typically packaged as command-line programs that expect to receive three main arguments: the path to the indexed reference sequence, the path to the input read file in FASTQ format, and the path to the output file in SAM or BAM format. This simple interface makes a number of assumptions: that all the required data files (reference, input and output) are accessible on locally mounted file systems; that the data are in the formats that the aligner supports; and that the use case supports executing a new process each time an alignment is required. Although these assumptions may appear to be reasonable within the conventional computing environment that is currently predominant in bioinformatics, they are actually extremely limiting to work aimed at introducing novel computing and data flow paradigms to the field – for instance, using distributed computing to improve scalability.

As a more general solution, we have defined a read aligner API that can be implemented with any underlying read mapping technology: the *Read Aligner Application Programming Interface (RAPI).^2^* For maximum compatibility the RAPI is defined in C. It includes generic functions and data structures to support typical alignment operations: index a reference sequence, load and unload the reference, map reads to the reference, interpret the results. Further, RAPI includes Python bindings, making it simple to load an aligner as a Python module and use it in unconventional ways – e.g., for scripting or in an interactive session. The project includes a reference aligner interface implementation that wraps the BWA-MEM aligner. This aligner plug-in, through its Python bindings, is used to compute alignments in Seal seqal. The fact that RAPI does not make any assumptions about the source or destination of the data makes it possible to easily integrate it with unconventional scalable computing and data storage technology, as has been done as part of this work. It makes it equally feasible to transparently implement aligner plug-ins based on GPGPU or FPGA accelerators. Also, since RAPI does not make any assumptions about the data formats, it also facilitates research into alternative data structures for persistent storage. Finally, since RAPI standardizes the aligner interface, one could parametrize the aligner to use within a RAPI-based pipeline, swapping aligner without changing any of the code.

RAPI is being proposed to the community as an option to standardize the read aligner interface. A standard interface would open up new use cases, reducing maintenance for existing applications, and make it simple and safe to harness aligner plug-ins to prototype and create novel functionality. The interface has been released under an open source license.

## VI. Evaluation

To evaluate the speed and scalability of our YARN-based sequence processing workflow we ran it on a real human sequencing dataset, with a varying number of nodes, processing the raw data in BCL format produced by the sequencers, reconstructing the DNA reads and aligning them to a reference genome. In the following subsections we describe all aspects of the experimental procedure and we compare it to a realistic baseline workflow.

### A. Hardware

All experiments were run on the Amazon Elastic Compute Cloud (EC2 – https://aws.amazon.com/ec2), using up to 16 instances of type r3.8xlarge. The specifications are provided in (Table II. Of note, the instances include *enhanced networking* which allows nodes to use Single Root I/O Virtualization, providing “higher I/O performance and lower CPU utilization, compared to traditional implementations.”^3^ Also, while the Intel CPUs provided by the instance provide 20 virtual cores each, the instance only has 32; we suppose that the instances run on dual-CPU hardware and the virtualization scheme hides 8 cores from the virtual machine.

**TABLE II:** Configuration of the Amazon EC2 r3.8xlarge nodes used to evaluate performance.

|  |  |
| --- | --- |
| **CPUs** | 32 virtual cores (Xeon E5-2670 v2, 25 MB cache) |
| **RAM** | 244 GiB |
| **Disks** | 2x320 GB SSD |
| **Network** | 10 Gb Ethernet connection with “enhanced networking” |
| **OS** | Debian 8 (Linux kernel 3.16) |

Finally, an Amazon Elastic Block Store (EBS) device was used as persistent storage for the input dataset, to avoid having to transfer the data to the compute center multiple times. However, this device was not used for I/O during any of our experiments.

### B. Input dataset

The input dataset is the output of a multiplexed sequencing run by an Illumina HiSeq 3000 at the CRS4 Sequencing and Genotyping Platform (at the same research organization as the authors of this manuscript). The run used a single multiplexing tag per fragment (the sequencing technology supports up to two) and contained 48 human genomic samples. The dataset contains 279 GB of raw data, of which 251 GB are read data scattered among 188,160 gzip-compressed BCL files (8 lanes × 112 tiles/lane × 210 cycles) plus 890 filter files (which contain a QC pass/fail for each read). The size of each uncompressed BCL and filter file is 4.1 MB. Additional information is reported in (Table III. Note that the input BCL files are automatically gzip-compressed by the software producing them on the sequencer.

**TABLE III:** Characteristics of the input dataset used for evaluation.

|  |  |
| --- | --- |
| **Sequencer** | Illumina HiSeq 3000 |
| **Total size of directory** | 279 GB |
| **Size of data files** | 251 GB (gzip-compressed) |
| **Number of files** | 1164005 |
| **Number of samples** | 48 |
| **Number of reads** | $2.9 \cdot 10^9$ |
| **Number of DNA bases** | $2.9 \cdot 10^{11}$ |

### C. Experimental Workflow

To evaluate the speed and scalability of our distributed-memory workflow, we prepared the following setup:

- HDFS distributed among the *n* computing nodes, with each datanode using its two SSD disks to store the data;
- YARN running on the same *n* nodes;
- YARN configured with 31 cores/node, 225 GB RAM/node, capacity scheduler with `DominantResourceCalculator`;
- one of the *n* nodes works as both master and slave, running the HDFS namenode, the YARN resource manager as well as the datanode and node manager services.

The input data are copied to the HDFS volume, along with a tar archive of the indexed reference required by Seal seqal for the alignment. Prior to each workflow invocation, the various system caches were cleared with `echo 3 > /proc/sys/vm/drop_caches`.

The experimental workflow then consists in running our Flink-based BCL conversion tool immediately followed by invoking Seal seqal for each resulting sample dataset.

#### 1) BCL conversion

To run our Flink-based BCL converter and demultiplexer, we start a detached Flink session on YARN. The session is configured with two TaskManagers per node, each with 80 GB of memory and 8 slots, while the Flink JobManager is assigned 10 GB of memory.

After the session is launched the BCL converter reads and write from/to HDFS. As soon as it completes its work, the Flink session on YARN is torn down by killing it (`yarn application–kill <appID>`). The operation produces PRQ files^4^ (a text-based, tab-separated, format that keeps paired reads in the same record) that are gzip-compressed at compression level 1.

#### 2) Alignment

Following the conclusion of the BCL conversion phase, its output directory is scanned to find the datasets produced. Empty files (as in gzip files with no content) are deleted to work around a bug in the InputFormat used by Seal seqal. The *Undetermined* dataset is also eliminated: these are the reads whose identifying tag was not recognized and thus cannot be assigned to any biological sample. Then, for each dataset, Seal seqal is invoked. All invocations are done in rapid succession, thus causing the various application instances to queue in YARN, which takes care to dispatch the numerous seqal tasks as computing resources become available. Seqal is configured to use 8 threads per task within the alignment core. On the other hand, the job is configured to request only two YARN cores per task; the effect is that in our experimental setup, with 31 cores per node, up to 15 tasks can run simultaneously (31/2 = 15). Seal seqal distributes the reference index to the various nodes via the Hadoop MapReduce distributed cache. This operation implies a small delay at the start of the first alignment tasks, as the archive is copied to all the nodes, unarchived, and the reference files are mapped to memory; however, subsequent tasks will find the reference already in memory and will benefit from caching effects. The seqal application directly reads the gzipped PRQ files produced by the BCL converter, processing one file per map task (gzip files are not splittable). It produces aligned reads in the SAM format [35] – a common text-based format for aligned reads – which are written to HDFS.

We ran this workflow on YARN clusters of 1, 2, 4, 8, and 16 nodes. On repeated runs we found very good agreement (within 2% of the mean running time). In the experiments we recorded the wall clock time required to run the entire procedure, starting at the launch of the Flink session on YARN and ending with the conclusion of the last alignment run; an intermediate time is also recorded at the end of the BCL conversion.

**TABLE IV:**
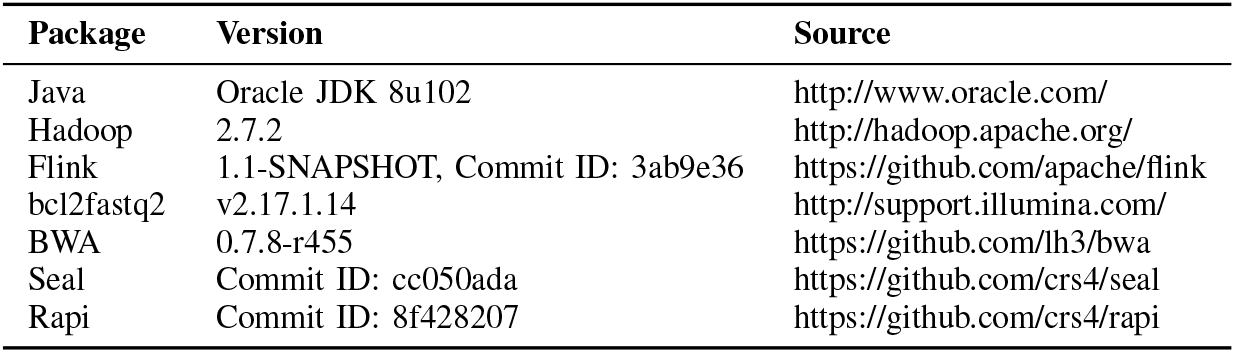
Versions of the various software packages used in this work.

### D. Baseline workflow

As the base case against which to compare, we implemented a workflow to perform BCL conversion and read alignment using the conventional tools described in Subsection II-C, thus using shared-memory parallelism on a single node. Therefore, the baseline workflow runs Illumina bcl2fastq2 and BWA-MEM (vers. 0.7.8-r455 – same version that is used by Seal seqal)

To prepare for the runs, the input data and the indexed reference is copied to one of the node’s SSD drives and the system caches are cleared in the same manner as described in Subsection VI-C.

The workflow is then started with the bcl2fastq2 tool. To allow it to fully exploit its multi-thread parallelism and minimize I/O latencies, we configured it to read BCLs from one SSD disk and write FASTQs to the other SSD disk. To lower its CPU usage we decreased the level of the output compression, by setting its option `--fastq-compression-level` to one (equivalent to gzip’s `--fast` option and identical to the setting used in our Flink-based converter). The tool detects the number of cores in the system and automatically selects the number of input, output, and processing threads to use; its settings appeared to be effective since it maintained a very high CPU utilization throughout the runs (very close to 100%).

Analogously to the YARN-based workflow, at the conclusion of the BCL conversion and demultiplexing step we remove the *Undetermined* dataset. The alignment is then performed with BWA-MEM. We configured the aligner to use 31 threads for its parallelized sections of code (option–t 31), which is analogous to YARN’s 31 cores/node. Since BWA-MEM does not natively read gzip-compressed files (and bcl2fastq2 does not write uncompressed files), we implemented the workflow to uncompress the files with gunzip on-the-fly and piped the data directly to the aligner, using process substitution and named pipes. On the other hand, BWA’s output, in SAM format just like Seal seqal’s, was redirected to the second SSD drive – i.e., BWA was reading from one drive and writing to the other.

Wall clock time was recorded at the start of the workflow, at the conclusion of bcl2fastq2, and at the end of the workflow.

## VII. Results & Discussion

### A. Full pipeline

The times recorded for the runs are shown in (Table V. The first, somewhat surprising, observation is that our suite is faster than the baseline even on a single node (618 vs 713 minutes). There are several explanations for this result:

1) Though the standard BCL to FASTQ conversion does take the same time on a single node (see VII-B), we optimized our pipeline using Seal’s PRQ file format which groups paired reads in the same file, thus resulting in half the files and less output data being written (perhaps ≈15%).
2) The BWA aligner used in the baseline processes reads in batches, computing alignments and then writing the results. Only the alignment phase seems to be parallelized, resulting in low CPU usage every time the aligner is generating output. On the other hand, our aligner runs more jobs concurrently, thus achieves a more uniform CPU usage while using the same BWA alignment core. This effect is illustrated in Fig. 3.
3) Seal `mmap’s` the reference data (thanks to our patch to the BWA core), meaning that after the first invocation seqal re-maps the reference in a tenth of a second. On the other hand, BWA `fread’s` the reference data at every invocation, which takes significantly longer and causes it to waste a significant amount of time over the 48 invocations of the experiment.
4) Our pipeline has better I/O characteristics than the baseline, since its tens of concurrent read and write streams (a variable number for Flink, while the aligner in this configuration had 31 read streams and 31 write streams), distributed uniformly over the two available SSD’s (through HDFS). The baseline, on the other hand, has four read and four write threads in bcl2fastq2 and two read and one write streams in the alignment phase. Moreover, all the reads are from one disk and all the writes are to the other disk.

**TABLE V:** Running times of our Hadoop/YARN pipeline and the baseline on a single node.

| Nodes | Time (minutes) |  |  |
| --- | --- | --- | --- |
|  | BCL | Align | Total |
| baseline | 67.3 | 645 | 713 |
| 1 | 47.3 | 571 | 618 |
| 2 | 24.5 | 286 | 311 |
| 4 | 13.5 | 144 | 158 |
| 8 | 8.1 | 72.5 | 80.7 |
| 16 | 6.1 | 36.6 | 42.8 |

Moving on to the multi-node results, given that both conversion and alignment can be decomposed in a large number of independent subtasks, we have been able to obtain an almost linear scaling, as shown in Fig. 2. We have tested strong scalability (i.e., increasing nodes while keeping the same input size) on up to 16 nodes and, thanks to our good performance on the single node test, our solution’s test result lies above the “perfect scalability” line of the baseline.

**Fig. 2:**
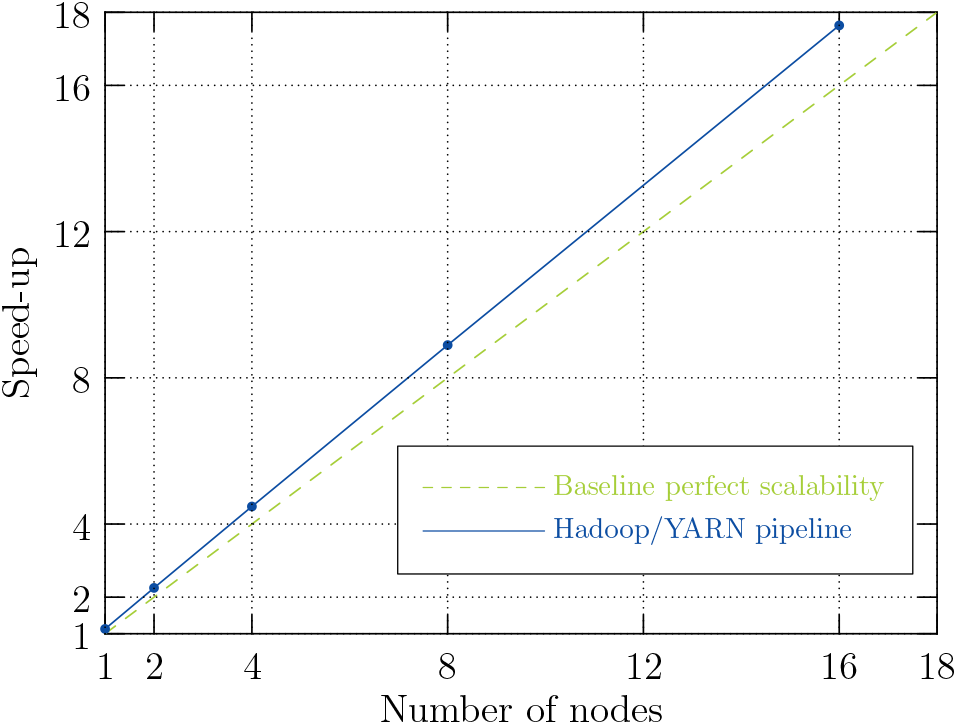
Strong scaling of our Hadoop/YARN pipeline, compared with the single-node baseline.

**Fig. 3:**
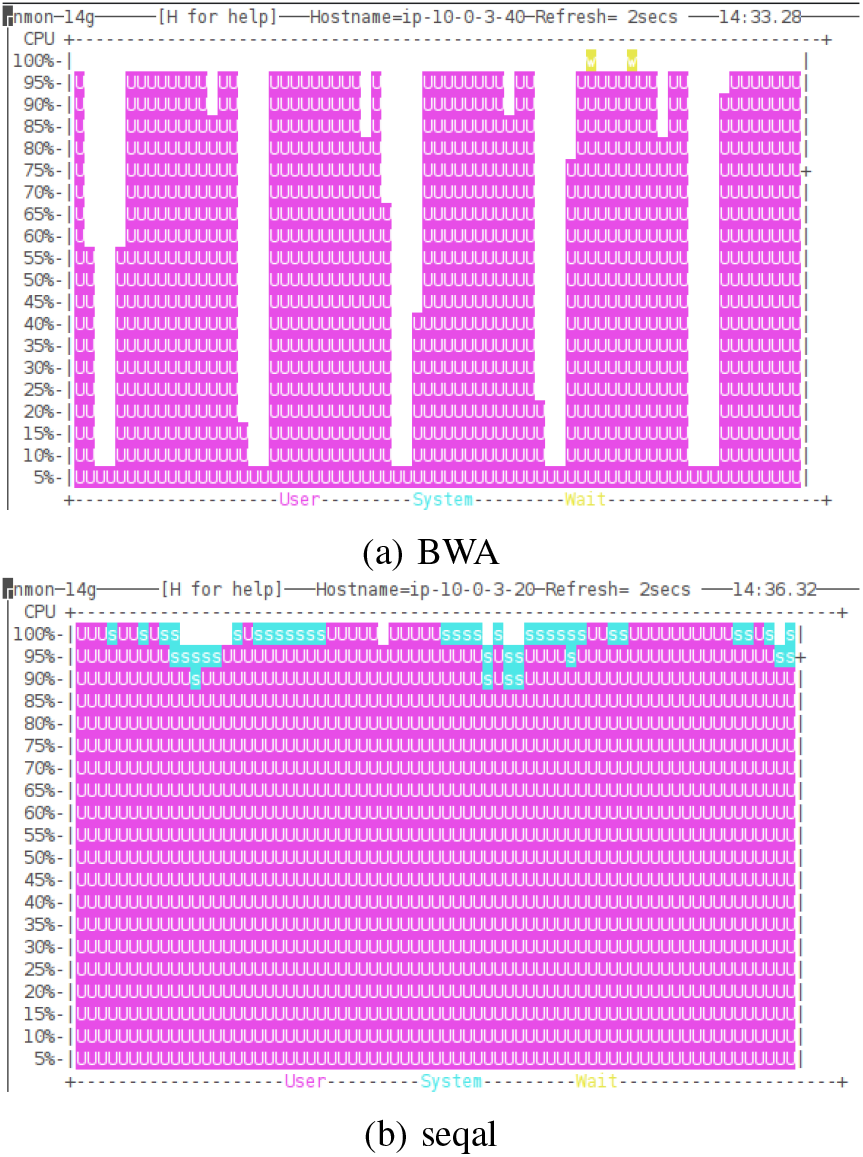
Computation intensity during alignment (`nmon` screenshots, *x* is time, *y* is CPU usage).

### B. BCL conversion

We also ran an independent test on our BCL conversion tool. To compare the performance of our core with Illumina’s bcl2fastq2 we implemented the gzip-compressed FASTQ output format, producing exactly the same output as Illumina’s tool. The results obtained in this test are shown in (Table VI and the strong, absolute scaling, compared to bcl2fastq2, is shown in (Fig. 4. We can see that on the single node we nearly match the performance of Illumina’s tool (58.4 vs 57.1 minutes) and we observe good scalability, which tapers as the running time goes below 10 minutes because of the fixed costs incurred by Flink’s scheduler. We have run this test on up to 14 nodes, choosing to stop at 14 nodes rather than 16 to have a more even division of the 896 = 64 · 14 Flink jobs.

**TABLE VI:** Running times of our Flink-powered BCL con verter with gzipped FASTQ output. Compare with Illumina bcl2fastq2: 57.1 minutes on a single node.

| Nodes | Time (minutes) |
| --- | --- |
| 1 | 58.4 |
| 2 | 30.2 |
| 4 | 15.2 |
| 6 | 10.5 |
| 8 | 8.4 |
| 14 | 5.4 |

**Fig. 4:**
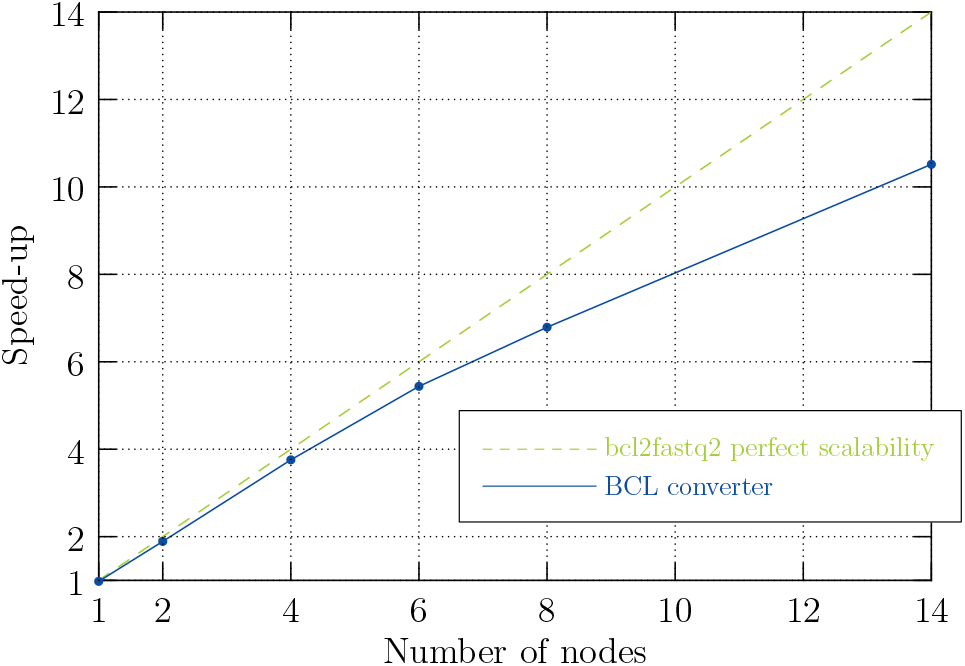
Speed-up of the Flink-powered BCL converter compared with the single-node Illumina bcl2fastq2 tool

The conversion from BCL to FASTQ handles as many data as the alignment, while taking much less time, because of its simpler algorithm. It is therefore the core which stresses more the I/O, making the analysis of the I/O performance in this phase a particularly interesting exercise.

Despite its relative simplicity the converter shows clearly to be CPU-bound, with all the virtual cores of the cluster at close to 100% utilization for the entire duration of the run. On a single node we observe a running time of 58.4 minutes, yielding a throughput of about 150 MB/s, considering both input and output, which is in fact much lower than the 700 MB/s that we could have theoretically extracted from the two SSDs of the machine, had we been limited by the I/O. When using 14 nodes the running time is only 5.43 minutes, implying a throughput of 1.6 GB/s during the program execution, which is particularly impressive considered that both input and output are compressed.

## VIII. Related work

Usage of “BigData” tools for bioinformatics applications started in late 2008 with work on Hadoop [36], [37] and has since continued to increase [38]. Most of the advancement in this area have come in NGS data analysis, where many Hadoop-based tools have been developed [39]; tools such as Crossbow [31] for SNP discovery, and Myrna [33] for RNA-Seq [40] differential expression analysis were pioneers in this area.

Several Hadoop-based sequencing alignment tools have already been published. Crossbow [31] (based on the Bowtie [41] aligner, which it executes internally) and Cloudburst [42] were two of the first. The alignment algorithms used by these programs has been superseded by others, including the BWA-MEM algorithm [17] which is used by Seal. More recently, newer distributed programs integrating BWA-MEM have been published, running on Hadoop MapReduce [29] and Spark [30], [43]. The authors have showed their tools to be faster than Seal’s latest official release, which is now obsolete because it integrates an outdated version of BWA. The version of Seal seqal used in this work is newer and though it has not been officially released it is open source and available on `github.com` in the `develop` branch of the project repository. Further, those works integrate BWA by running the executable from within their framework; Seal uses a more flexible approach which will more easily allow future work in the area of integrating more advanced storage file formats into the NGS processing workflows.

Of course, there exist other approaches to scale the throughput of sequence data processing. Work has been done on using hardware “accelerators” such as Graphical Processor Units (GPUs) [44], [45] or Field-Programmable Gate Arrays (FPGAs) [46]–[48] to accelerate sequence alignment. These should be seen as complementary to our work since they could be easily integrated into our Seqal aligner by implementing a RAPI-compatible plugin.

It is also worth mentioning a project called ADAM [49] that is developing a suite of tools for the analysis of sequencing data on Spark. That project is also complementary to this work since it provides a myriad of tools that are useful for post-alignment sequence analysis.

Finally, an important project relevant to this work is the Genome Analysis Toolkit (GATK) [50], which implements essential analyses for NGS data that are downstream of alignment. Moreover, the GATK is probably one of the most important packages in its niche, and independent research has found it to be one of the best at what it does [18], [19]. The new version of the GATK will run on Spark, which makes it an excellent complement for the work presented in this manuscript: with our YARN-based pipeline and GATK 4 a sequencing center could implement the entire processing pipeline on YARN.

## IX. Conclusion

We have presented a YARN-based pipeline to process raw Illumina NGS data up to the stage of aligning reads – the first, to the best of the authors’ knowledge, to process raw data. Our experiments have shown the pipeline to have excellent scalability characteristics, such that a sequencing center could reasonably aim to reduce their processing per sequencing run to under an hour with the use of a small YARN cluster. Moreover, our solution performed better than the baseline even on a single computing node. Completing this pipeline has required the creation of a Read Aligner API and improvements to the efficiency of the sequence aligner in the Seal suite of tools. The work we presented is an excellent complement to work currently being done by the GATK group to bring the sequence analysis downstream of alignment to the YARN platform; combining our tools one could have a complete YARN-based pipeline for NGS data, and then further improve performance by adopting an in-memory file system such as Apache Arrow, thus removing the need to write intermediate data to disk.

The code presented it this work is already available as open source.^5^

## Acknowledgments

We thank Gianmauro Cuccuru for the NGS dataset used in the experiments and Massimo Gaggero for his support in setting up the AWS EC2 environment. Luca Pireddu would also like to thank Roman Valls Guimera for his useful suggestions and insights.

## Footnotes

1 An updated list of users is available at: https://cwiki.apache.org/confluence/display/FLINK/Powered+by+Flink

2 https://github.com/crs4/rapi

3 https://aws.amazon.com/ec2/faqs/#What_networking_capabilities_are_included_in_this_feature

4 http://biodoop-seal.sf.net/file_formats.html#prq-file-format

5 Code available at http://github.com/crs4

